# Re-visiting the potential impact of doxycycline post-exposure prophylaxis (doxy-PEP) on the selection of doxycycline resistance in *Neisseria* commensals

**DOI:** 10.1101/2025.01.09.632169

**Authors:** Leah R. Robinson, Caroline J. McDevitt, Molly R. Regan, Sophie L. Quail, Makenna Swartz, Crista B. Wadsworth

## Abstract

Doxycycline post-exposure prophylaxis (doxy-PEP) is a preventative strategy demonstrated to reduce bacterial sexually transmitted infections in high-risk populations. However, the impact of doxy-PEP on antibiotic resistance acquisition in key members of our microbiomes, is as of yet unclear. For example, commensal *Neisseria* are known reservoirs of resistance for gonococci through horizontal gene transfer (HGT), and are more likely to experience bystander selection due to doxy-PEP as they are universally carried. Thus, the consequences of doxycycline selection on commensal *Neisseria* will be critical to investigate to understand possible resistance mechanisms that may be transferred to an important human pathogen. Here, we use *in vitro* antibiotic gradients to evolve four *Neisseria* commensals (*N. cinerea, N. canis, N. elongata, and N. subflava,* n=4 per species) across a 20-day time course; and use whole genome sequencing to nominate derived mutations. After selection, 12 of 16 replicates evolved doxycycline resistance (> 1 μg/mL). Across resistant lineages: An A46T substitution in the repressor of the Mtr efflux pump (MtrR) and a V57M substitution in the 30 ribosomal protein S10 were clearly associated with elevated MICs. Additional mutations in ribosomal components also emerged in strains with high MICs (i.e., *16S rRNA G1057C*, RplX A14T). We find the MtrR 46T, RpsJ 57M, and RplX 14T circulating in natural commensal populations. Furthermore, *in vitro* co-evolution of *N. gonorrhoeae* with *Neisseria* commensals demonstrated rapid transfer of the pConj plasmid to *N. subflava* and *N. cinerea*, and p*bla* to *N. cinerea*. Finally, collection of novel commensals from human hosts reveals 46% of isolates carrying doxycycline resistance; and doxycycline resistance was significantly greater in participants self-reporting doxycycline use in the past 6 months. High-level doxycycline resistance (> 8 μg/mL) was always associated with carriage of the ribosomal protection protein (*tetM)* and pConj. Ultimately, characterizing the contemporary prevalence of doxycycline resistance, and underlying resistance mechanisms, in commensal communities may help us to predict the long-term impact of doxy-PEP on *Neisseria*, and the likelihood of transferring particular genotypes across species’ boundaries.

## Introduction

The use of the tetracycline-class antibiotic doxycycline as a post-exposure prophylaxis treatment (200 mg of doxycycline 24-72 hours after unprotected intercourse), commonly referred to as doxy-PEP, has been recently recommended by the CDC and other public health organizations to prevent the transmission of sexually transmitted infections (STIs) (i.e., gonorrhea, chlamydia, and syphilis) (Bachmann *et al*. 2024). Doxycycline is a broad-spectrum antibiotic that inhibits protein production by binding allosterically to the 30S ribosomal subunit of bacterial ribosomes and preventing the association of aminoacyl-tRNA with the bacterial ribosome However, due to the high carriage rate of conjugative plasmids (pConj) containing the ribosomal protection protein (*tetM)* in gonococcal populations (e.g., ∼33% in the United States, and up to 90% in LMICs (Cehovin and Lewis 2017; Cehovin *et al*. 2020)) doxycycline is not recommended to treat gonococcal infections. Doxy-PEP was trialed in 2015-2016 (Molina *et al*. 2018) and 2020-2022 (Luetkemeyer *et al*. 2023b) with some success, however there is concern among the members of the *Neisseria* research community that doxy-PEP will select for a higher prevalence of both doxycycline resistant and multidrug resistant gonococci (MDR-Ngo) (Kong, Kenyon and Unemo 2023; Reichert and Grad 2024; Vanbaelen, Manoharan-Basil and Kenyon 2024). In the United States, doxy-PEP has been shown to decrease the rate of acquiring STIs by 20% compared to standard care (Luetkemeyer *et al*. 2023b). This study also showed a decreased incidence of gonorrhea (reduced by 10%), though higher rates of tetracycline resistant *N. gonorrhoeae* in doxy-PEP participants. In France (the IPERGAY study), doxy-PEP reduced the risk of all bacterial STIs in high-risk men who have sex with men (MSM) populations, except for gonococcal infections (Molina *et al*. 2018); with high rates of tetracycline resistant gonococci in Europe (average = 63.4%; France = 92.3%) likely contributing to this outcome (Unemo *et al*. 2024). Mathematical modeling predicts that doxy-PEP will reduce gonococcal infections in the short-term as susceptible lineages are killed, however will increase the burden of gonorrhea long-term as doxycycline resistant lineages spread (Reichert and Grad 2024). These models also predicted reduced clinical utility of doxycycline within 10-20 years, and little impact on extending the usefulness of ceftriaxone as a treatment for gonorrhea (Reichert and Grad 2024). *In silico* analysis suggest even that doxy-PEP may select for cross-resistance to ceftriaxone in *N. gonorrhoeae* (Vanbaelen, Manoharan-Basil and Kenyon 2023; Whiley *et al*. 2023).

Doxy-PEP may also have unintended consequences within the commensal *Neisseria* as a result of bystander selection. Given that commensal *Neisseria* have a higher carriage rate (100%) compared to the pathogenic *Neisseria* (0.01% to 10%) (Diallo *et al*. 2016; Kenyon and Schwartz 2018; Dong *et al*. 2020; Vanbaelen *et al*. 2022), their exposure to unintentional antibiotic selection is higher. Human-associated *Neisseria* commensals colonize the naso- and oropharyngeal niche and have transferred important resistance phenotypes to the pathogenic *Neisseria* through horizontal gene transfer (HGT) (Goytia and Wadsworth 2022). For example, β-lactam (through the acquisition of commenal *penA* alleles (Spratt, Brian 1988; Ameyama *et al*. 2002)), macrolide (*mtr* (Rouquette-Loughlin *et al*. 2018; Wadsworth *et al*. 2018)), and quinolone resistance (*gyrA* (Gorla *et al*. 2018; Chen *et al*. 2020)) have all been passed from commensals to pathogens; indicating the importance of these species as reservoirs of resistance. Doxycycline resistance in *Neisseria* commensals has been reported to be high (∼63%), and though doxy-PEP was only marginally found to increase doxycycline resistance in commensal *Neisseria* (63% resistant at baseline compared to 69% at month 12 post doxy-PEP, *p* = 0.11) (Luetkemeyer *et al*. 2023a), highlights the selective potential of doxy-PEP on driving resistance emergence in commensal communities. Thus, it is critical to characterize the identity of doxycycline resistance mechanisms in commensal *Neisseria* species in order to predict genotypes that may ultimately be transferable to pathogenic *Neisseria* species. There is a single study that has attempted to characterize doxycycline resistance mutations in *Neisseria* commensals emerging post-selection in the laboratory, however there was no evidence of resistance emergence after seven days of drug exposure (for *N. gonorrhoeae* and *N. subflava*) (Kenyon *et al*. 2023). This limited focus on doxycycline resistance in commensals leaves a ‘black box’ as to which commensal resistance mutations may spread as a result of doxy-PEP selection, both within commensal communities and perhaps to the pathogenic *Neisseria* as well. Our understanding of doxycycline resistance in *Neisseria* is mostly limited to the pathogenic *Neisseria*, and is mediated by both plasmid-borne and chromosomal mutations. Plasmid-mediated doxycycline resistance is harbored on the pConj plasmid, which both enables conjugation and contains the *tetM* gene, encoding a ribosomal protection protein (Morse *et al*. 1986; Pachulec and Van Der Does 2010; Cehovin and Lewis 2017; Cehovin *et al*. 2020). There are three main types of gonococcal pConj plasmids: A 24.5 MDa plasmid (pLE2451, with three different variants: pConj.5, pConj.6, and pConj.7), and two 25.2 MDa plasmids (Dutch-type (two variants: pConj.3 and pConj.4) and American-type (two variants: pConj.1 and pConj.2)) containing the *tetM* gene. The *tetM* determinant on gonococcal 25.2 MDa pConj plasmids originated from a class II Tn916-like streptococcal transposon, which has had the transposase deleted while maintaining the functionality of the *tetM*-region (Swartley *et al*. 1993). Concerningly, pConj plasmids have been shown to be transferred from *N. gonorrhoeae* to the commensal *Neisseria* and back again to *N. gonorrhoeae* in the laboratory (Genco, Knapp and Clark 1984; Roberts and Knapp 1988; Roberts and Knapp 1988), and have also been identified in natural commensal populations (*N. meningitidis*, *N. subflava, N.lactamica,* and *N. polysaccharea*) (Yee *et al*. 2023); suggesting cross-species barriers to plasmid transfer, and high MIC tetracycline and doxycycline resistance, are limited. Additionally, the main chromosomal mutation contributing to tetracycline-class antibiotic resistance in gonococci is a RpsJ (ribosomal S10 protein) V57M substitution located proximal to the aminoacyl-tRNA site in the 30S ribosomal subunit, altering the structure near the tetracycline-binding site, and decreasing antibiotic affinity (Hu *et al*. 2005). However, mutations in PorB, MtrR, and PilQ have also shown some impact on tetracycline MICs (Hu *et al*. 2005). As in our prior work (Raisman *et al*. 2022; Frost *et al*. 2024), it will be important to enumerate the paths to doxycycline resistance in diverse commensal isolates to identify if resistance is conserved across the genus, or if different adaptive solutions exist.

Though tetM-containing pConj plasmids have already been identified in *Neisseria* commensals (Yee *et al*. 2023), contributing chromosomal resistance mutations in commensals are unclear (Fiore *et al*. 2020); and given the prediction that doxy-PEP will increase doxycycline resistant gonococcal lineages (Reichert and Grad 2024), we anticipate doxycycline resistant commensal lineages will also increase; therefore, their underlying resistance mechanisms will be critical to characterize. Thus, we set out to characterize doxycycline resistance in *Neisseria* commensals using a multi-faceted approach. *In vitro* culture using gradient-based selection with several commensal *Neisseria* has been used to flesh out the *Neisseria* resistome for azithromycin, ciprofloxacin, and penicillin previously (Raisman *et al*. 2022; Frost *et al*. 2024; Robinson *et al*. 2024). Here, we expand this method to explore doxycycline resistance-emergence in the commensal *Neisseria*; and query the presence of *in vitro* derived mutations in natural *Neisseria* populations. We also examine the potential for cross-species’ plasmid transfer of pConj and the other gonococcal plasmids in co-culture. Finally, we collect a novel panel of *Neisseria* commensal isolates and assess the prevalence of doxycycline resistance in those isolates, the impact of doxycycline use on the emergence of resistance, and doxycycline resistance mutations within novel isolates.

## Materials and Methods

### Bacterial strains and culture conditions

This study utilized the bacterial strains AR-0944 (*N. cinerea*), AR-0945 (*N. elongata*), AR-0948 (*N. canis*), and AR-0953 (*N. subflava*), all acquired from the Centers for Disease Control and Prevention (CDC) and Food and Drug Association’s (FDA) Antibiotic Resistance (AR) Isolate Bank “*Neisseria* species MALDI-TOF Verification panel”. All selected commensals were human-associated, except for *N. canis*, which colonizes the oral cavity of dogs and cats but has also been isolated from human patients with bite wounds (Bailie, Stowe and Schmitt 1978; Guibourdenche, Lambert and Riou 1989). All isolates had been previously sequenced (Fiore *et al*. 2020; Frost *et al*. 2024). Strains were cultured on GC agar base (Becton Dickinson Co., Franklin Lakes, NJ, USA) media plates containing 1% BBL IsoVitaleX Enrichment (Becton Dickinson Co.,Franklin Lakes, NJ, USA; hereafter, GCB-I media), and were grown for 18–24 hours at 30°C. Stocks were created in trypticase soy broth (TSB) containing 20% glycerol, and stored at −80°C.

### *In vitro* doxycycline selection

Doxycycline selection was conducted by passaging four replicates of each commensal species using a selective gradient of antibiotic created with an ETEST strip (Biomérieux, Durham, NC, USA) on GCB-I media. In brief, cells collected from above the zone of inhibition (ZOI) and a 1 cm band below the ZOI were suspended in TSB and spread onto new GCB-I plates, along with a 0.016-256 μg/mL doxycycline ETEST strip, and were incubated at 30°C. After 48 hours, MICs were recorded. This process was repeated for a total of 11 passages over the course of 20 days for each bacterial replicate. Final bacterial populations were struck onto GCB-I plates and individual colonies were grown and stocked for further analysis. Control strains were passaged in the absence of selection using the same protocol.

MICs for single colony stocks of post-selection lineages were measured via Etest strips, which have been demonstrated to have comparable MIC values to the agar dilution method (Papp *et al*. 2018). In brief, cells from overnight plates were suspended in TSB to a 0.5 McFarland standard and inoculated onto a GCB-I plate with an Etest strip. Following 18–24 hours of incubation at 30°C, the MICs of each replicate were recorded. We defined resistance as an MIC ≥ 2 μg/mL, the Clinical Laboratory Standards Institute (CLSI) breakpoint for tetracycline for *N. gonorrhoeae* (Clinical and Laboratory Standards Institute 2020). Reported MICs are the mode of 2-3 independent tests and were agreed upon by two independent researchers.

### Whole genome sequencing and bioinformatics pipeline

Cell lysis was performed by suspending cell growth from overnight plates in TE buffer (10 mM Tris [pH 8.0], 10 mM EDTA) with 0.5 mg/mL lysozyme and 3 mg/mL proteinase K (Sigma-Aldrich Corp., St. Louis, MO). DNA isolation utilized the PureLink Genomic DNA Mini Kit (Thermo Fisher Corp., Waltham, MA) with RNase A treatment to remove RNA. Isolated DNA was prepared for sequencing using the Nextera XT kit (Illumina Corp., San Diego, CA), and uniquely dual-indexed and pooled. The final pool was subsequently sequenced on the Illumina MiSeq platform at the Rochester Institute of Technology Genomics Core using V3 600 cycle cartridges (2×300bp). Quality of each paired-end read library was assessed using FastQC v0.11.9 (Andrews 2010). Adapter sequences and poor-quality sequences were trimmed from read libraries using Trimmomatic v0.39, using a phred quality score < 15 over a 4 bp sliding window as a cutoff (Bolger, Lohse and Usadel 2014). Reads < 36 bp long, or those missing a mate, were also removed from subsequent analysis. Reference genomes were assembled and annotated previously (Fiore *et al*. 2020; Frost *et al*. 2024), and read libraries were mapped back to these reference assemblies using Bowtie2 v.2.2.4 using the “end-to-end” and “very-sensitive” options (Langmead and Salzberg 2012). Finally, Pilon v.1.16 was used to identify derived mutations (Walker *et al*. 2014); and custom scripts were used to determine if they were located in annotated genomic features. SNPs present in control strains were omitted from subsequent consideration as resistance candidates. Amino acid positions were numbered according to a gonococcal reference strain (NZ_AP023069.1).

### Assessment of prevalence of derived mutations in natural populations

We used isolates deposited to the PubMLST database (https://www.pubmlst.org/Neisseria) (Jolley, Bray and Maiden 2018) to interrogate the presence of experimentally derived mutations in natural *Neisseria* populations, and to investigate the possibility of locus-level cross-species HGT. We downloaded 2116 alleles from PubMLST for *mtrR*, *rpsJ*, the *16S rRNA,* and *rplX* for all human-associated commensal *Neisseria* (*N. benedictiae, N. bergeri, N. blantyrii, N. cinerea, N. elongata, N. lactamica, N. maigaei, N. mucosa, N. oralis, N. polysaccharea, N. subflava, N. uirgultaei* and *N. viridiae)* (Diallo *et al*. 2019). *N. gonorrhoeae* was included as the focal pathogen in this analysis, however *N. meningitidis* was excluded. We limited the majority of the gonococcal dataset to the last five years (2020-2024) to capture more recent commensal-pathogen transfer events, though included commensals isolates from all years, and added targeted *N. gonorrhoeae* isolates of interest isolated from prior years. Alleles for each gene were aligned using MAFFT v.7 (Katoh and Standley 2013) with a maximum likelihood tree calculated using RAxML v.8.0 (Stamatakis 2014). iTOL was used to visualize and annotate the resultant trees (Letunic and Bork 2021).

### Pathogen and commensal short-term co-evolution experiment

The *N. gonorrhoeae* WHO N reference strain obtained from the Centers for Disease Control and Prevention (CDC) and Food and Drug Association’s (FDA) Antibiotic Resistance (AR) Isolate Bank “*Neisseria gonorrhoeae* WHO Reference” panel was used as a plasmid donor strain for co-culture experiments. WHO N harbors pConj and p*bla* plasmids (Unemo *et al*. 2016), which confer tetracycline/doxycycline and penicillin resistance respectively, as well as the cryptic plasmid. This strain has been phenotyped for its minimum inhibitory concentration (MIC) to these drugs previously (MIC to tetracycline = 16 μg/mL, MIC to penicillin > 32 μg/mL) (Unemo *et al*. 2016), which was confirmed in our laboratory.

We co-evolved WHO N with each commensal *Neisseria* species described above for 7 days on GCB-I plates at 37°C in a 5% CO_2_ incubator. Strains were first added to GCB-I plates at equivalent concentrations (100 ul of a 0.5 McFarland standard suspension of cells in TSB), and subsequently passaged to new GCB-I plates every 24 hours. After 7 days, the entire population was re-plated on GCB-I plates containing doxycycline (6 μg/mL). Single colonies with commensal morphology were selected (n=2 per species), struck on to new plates, propagated for another 24 hours, and then stored in trypticase soy broth (TSB) containing 20% glycerol at - 80°C for further analysis. Subsequently, from these single colony picks, MICs were determined for doxycycline and penicillin using Etest strips as described above. DNA was isolated, and whole genome sequencing was conducted as described above. After trimming read libraries with Trimmomatic v0.39 (Bolger, Lohse and Usadel 2014) using the aforementioned parameters; read libraries were assembled using SPAdes v.3.14.1 (Bankevich *et al*. 2012); and the presence of pConj, p*bla*, and pcryp was assessed in commensal draft assemblies using blastn (Camacho *et al*. 2009). Species identity for read libraries were confirmed using kraken2 (Wood, Lu and Langmead 2019).

### Characterization of doxycycline resistance in a novel panel of commensal *Neisseria* isolates

Study participants (≥ 18 years old, n=15) were recruited from students, staff, and faculty at the Rochester Institute of Technology (RIT) main campus (USA: Rochester, NY) in the fall of 2024. The study was approved by the Institutional Review Board of RIT (protocol number: 02030524, date of approval: 30 May 2024). Participants were provided written informed consent prior to participation in the study. After consent, participants were asked to complete a voluntary demographic questionnaire, including information on their: age, sex, and past antibiotic use over the past 6 months. For bacterial collection, participants agitated the surfaces of their mouths for 30 seconds with their tongue, and deposited their spit sample into a sterile container. Spit samples were subsequently inoculated onto LBVT.SNR agar (Knapp 1988; Knapp and Hook 1988), which promotes the growth of *Neisseria* commensals, and grown for 48 hours at 30°C. Ten colonies were selected from each participant for propagation and further analysis. For each, DNA was isolated, and whole genome sequencing was conducted as described above. However, these read libraries were sequenced on the NovaSeq 6000 platform at the RIT Genomics Core. After trimming read libraries with Trimmomatic v0.39 (Bolger, Lohse and Usadel 2014), species identity was confirmed using kraken2 (Wood, Lu and Langmead 2019). Reads were assembled with SPAdes v.3.14.1 (Bankevich *et al*. 2012) and blastn (Camacho *et al*. 2009) was used to confirm the presence of experimentally derived mutations, *tetM,* and β-lactamase genes. Identified *Neisseria* were stored in TSB and 20% glycerol at −80°C. Doxycycline MICs were recorded by suspending cells in TSB and spreading suspensions onto GCB-I plates, along with a 0.016-256 μg/mL doxycycline ETEST strip. Plates were incubated at 30°C for 24 hours and the mode of 2-3 tests was selected as the final MIC.

## Results

### Selection for elevated doxycycline MICs in commensal Neisseria

Four *Neisseria* species were passaged on doxycycline gradients to select for mutations imparting reduced susceptibility. The species included: *N. cinerea* [AR-0944], *N. subflava* [AR-0953], *N. elongata* [AR-0945], and *N. canis* [AR-0948]. We phenotyped ancestral non-selected strains for their doxycycline MICs which are displayed in Table 1, and ranged from 0.38-2 μg/mL. After 20 days of selection, selected populations of each species had at least one replicate evolving an elevated MIC compared to the ancestral strain (Figure 1). Single colonies from each selected population were selected and propagated for downstream analysis. Of these lineages grown from single colonies, 50% of *N. cinerea* isolates evolved elevated doxycycline MICs compared to the ancestral strain (mean = 3 ± 1.15); 50% of surviving *N. elongata* isolates evolved elevated MICs (mean = 0.75 ± 0.35); 100% of *N. canis* isolates evolved elevated MICs (mean = 6 ± 0); and 100% of *N. subflava* isolates evolved elevated MICs (mean = 8 ± 5.66) (Table 1).

**Figure 1.**
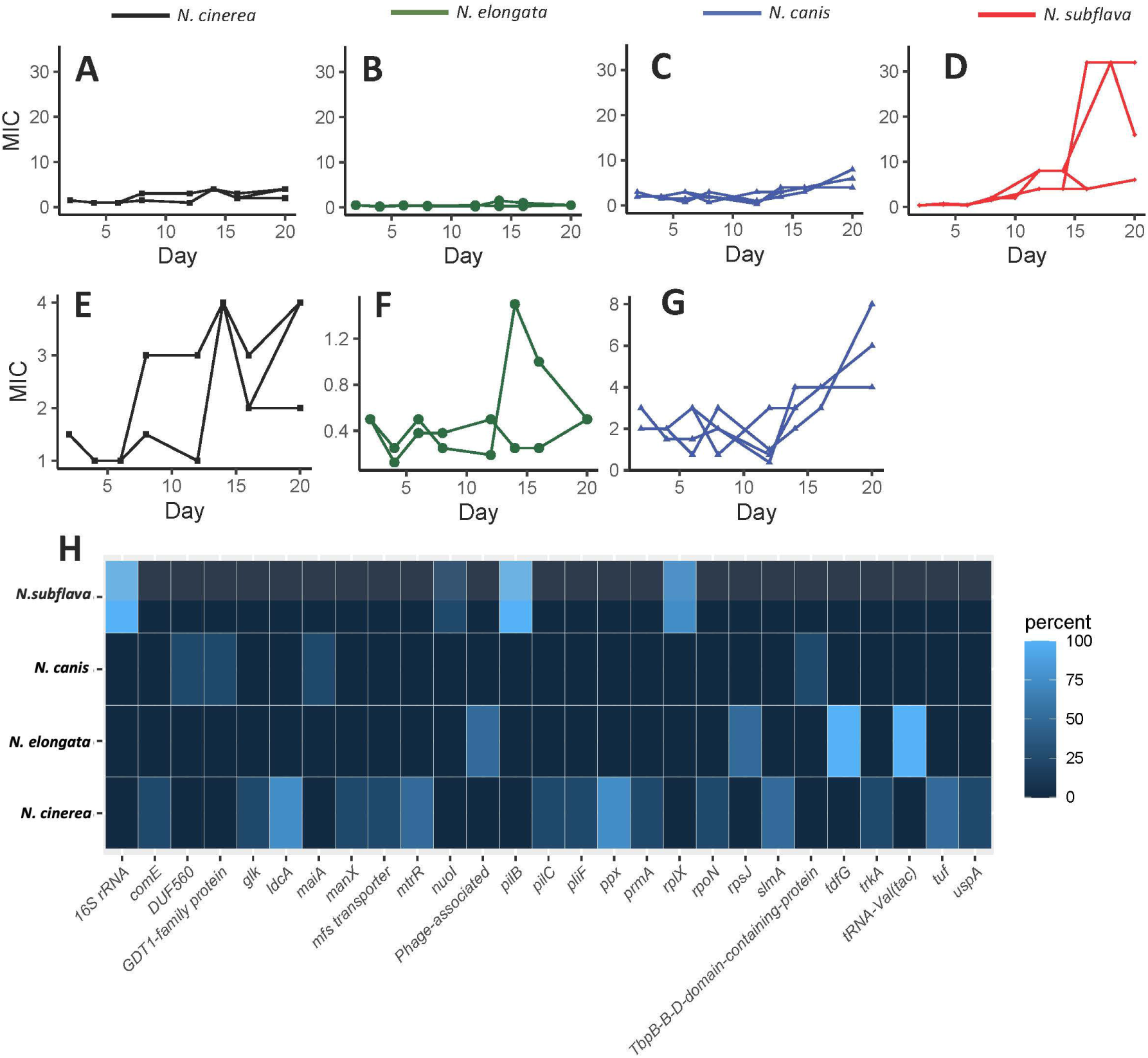
*In vitro* selection of four commensal Neisseria species with doxycycline. In brief, four experimental replicates of each species were passaged for 20 days on selective gradients created with Etest strips. Cells for each passage were selected by sweeping the zone of inhibition (ZOI) and a 1 cm band in the bacterial lawn surrounding the ZOI. MIC trajectories are shown for (A) *N. cinerea*, (B) *N. elongata*, (C) *N. canis*, and (D) *N. subflava*. Expanded scales are shown in panels E-G for *N. cinerea*, *N. elongata*, and *N. canis* respectively. (H) Coding domain (CD) identity of *in vitro* derived mutations for drug-selected lineages. The percentage of mutational hits within a given gene is displayed as a heatmap, with brighter blue coloration indicating more frequent occurrence of a mutation within a CDS in replicate evolved lineages for each species.

**Table 1.** Derived haplotypes for replicate experimental lineages post-selection.

| Species | Doxycycline MIC (µg/mL) |  |  |  |  |
| --- | --- | --- | --- | --- | --- |
|  | Ancestral | Replicate 1 | Replicate 2 | Replicate 3 | Replicate 4 |
| *N.cinerea* | 2 | 2 [*IdcA*, *ppx*, *prmA*] | 4 [**<u>MtrR A46T</u>**, **<u>pilF</u>**, **<u>trkA</u>**] | 2 [*tuf*, *IdcA*, *ppX*, *rpoN*] | 4 [**<u>comE</u>**, *tuf*, *IdcA*, **<u>MtrR A46T</u>**, *ppx*, *uspA*] |
| *N.elongata* | 0.5 | DNG | 0.5 [*tonB*] | DNG | 1 [*tonB*, **<u>RpsJ V57M</u>**] |
| *N.canis* | 2 | 6 [**<u>GDT1-family protein 226-CTACCA</u>**] | 6 [*maiA A484C-silent*] | 6 [**<u>TbpB-B-D domain protein duplication</u>**] | 6 [**<u>DUF560 domain protein duplication</u>**] |
| *N.subflava* | 0.38 | 8 [**<u>pilB-insertion, 16S rRNA G1057C</u>**] | 4 [**<u>pilB-insertion, RplX A14T, 16S rRNA G1057C</u>**] | 4 [*nuoI*, **<u>pilB-insertion, RplX A14T, 16S rRNA G1057C</u>**] | 16 [**<u>pilB-insertion, RplX A14T, 16S rRNA G1057C</u>**] |

DNG = Did not grow. [ ] genes within brackets indicate the specific loci with mutations post-selection in experimental lineages, **bolded** genes indicate those unique to strains with elevated MICs, **bolded and underlined** genes indicate consistently mutated loci associated with elevated MICs post-selection.

### Comparative genomics and nominating derived mutations

Whole genome sequencing was conducted for experimentally evolved lineages to nominate derived mutations post-selection. Mutations were mapped to replicate lineages, to assess which mutations were the most likely to be causal to elevated MICs (Table 1). For *N. cinerea*, mutations emerged in 14 loci (Figure 1), however only inheritance of a A46T substitution in the repressor of the Mtr efflux pump (MtrR) was consistently associated with elevated MICs in 2 out of 4 isolates. Strains inheriting these mutations had elevated doxycycline MICs of 4 μg/mL compared to the ancestral lineage with an MIC of 2 μg/mL. For *N. elongata*, mutations emerged in 4 loci (Figure 1), however only the 30 ribosomal protein S10 (encoded by *rpsJ*) mutation V57M was associated with elevated doxycycline MICs (1 μg/mL, compared to 0.5 μg/mL for the ancestral lineage). All *N. canis* isolates had elevated doxycycline MICs at 6 μg/mL compared to the ancestral of 2 μg/mL, yet they all inherited different mutations: One lineage had an in-frame insertion CTACCA at position 226 in a GDT1-family-protein, the second inherited a silent A to C substitution at position 484 in maleylacetoacetate isomerase (*maiA)*, the third a putative duplication of the gene encoding a TbpB-B-D domain containing protein, and the fourth a putative duplication of the gene encoding a DUF560 domain containing protein (Table 1). Finally for *N. subflava*, all lineages had elevated MICs ≥ 4 μg/mL; and all inherited a *16S ribosomal RNA* G to C substitution at position 1057 and an insertion in the type IV pilus assembly ATPase (*pilB*) altering reading frame; with 3 of 4 strains also inheriting several mutations in the 50S ribosomal protein L24 (*rplX*), however the only common amino acid-altering substitution was A14T (Table 1).

### Prevalence of MtrR, RpsJ, RplX, and 16s rRNA variants in natural *Neisseria* populations and locus-level HGT characterization

We searched for evidence of *in vitro* selected mutations and gene-level recombination tracts in natural *Neisseria* populations using a subset of *Neisseria* isolates deposited to the PubMLST database (n=2116). While the *16S rRNA* 1057 site was invariable in natural *Neisseria* populations, with no detected recombination events between *N. gonorrhoeae* and commensals, we did observe variation at other loci. For example, for *mtrR*, the amino acid-altering mutation A46T occurred in 2 of 4 *in vitro* selected *N. cinerea* isolates, and was consistently associated with elevated doxycycline MICs. In natural populations of *Neisseria*, we observe a diversity of alleles at *mtrR* encoding different amino acids with: 1929 isolates harboring an alanine (A), 92 with a threonine (T), 1 with a glutamic acid (E), and 81 with an arginine (R) (Figure 2). All observed threonines at position 46 occurred within *N. gonorrhoeae* isolates in the wild-type *mtrR* clade (Figure 2). Three *N. lactamica* isolates also harbored a threonine at position 46 however (37003 (isolated in the UK in 2015), 44098 (isolated in the UK in 2015), and 88815 (isolated in Brazil in 2014)). An additional large gonococcal clade was observed with more *N. lactamica and N. polysaccharea*-like alleles at *mtrR* (Figure 2).

**Figure 2.**
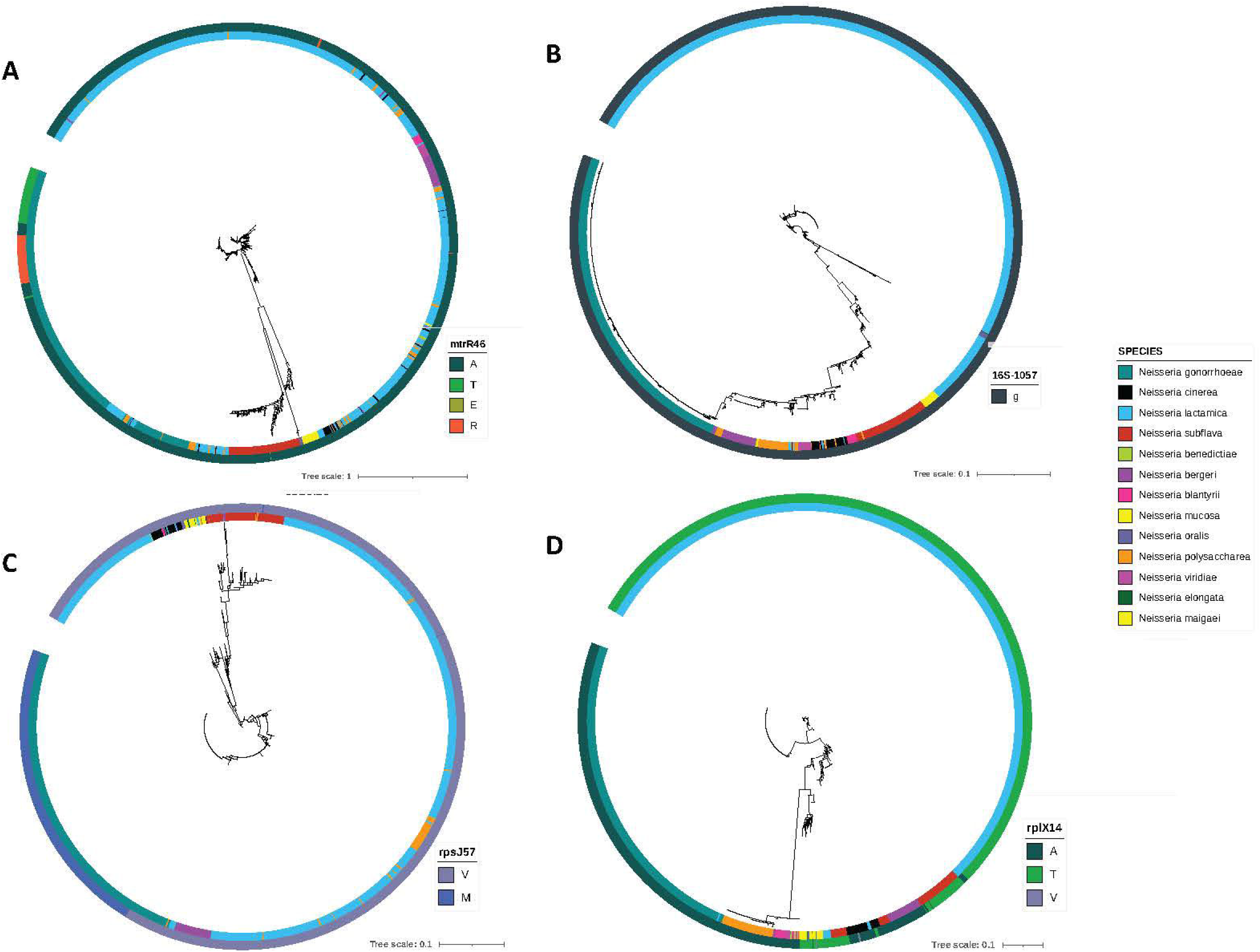
Characterizing experimentally derived mutations in natural *Neisseria* populations. Maximum likelihood phylogenies for *mtrR*, the *16S rRNA*, *rpsJ*, and *rplX* for 2116 PubMLST *Neisseria* isolates. (A) The derived MtrR A46T substitution was more prevalent in gonococci (17%) than commensals (0.1%). There are two clear gonococcal *mtr* clades, indicative of interspecies admixtures. (B) The *16S rRNA* 1057 site was invariable in natural *Neisseria* populations, with no detected recombination events. (C) The derived RpsJ V57M was rare in commensal isolates (∼0.3%) however had a high carriage rate in gonococci (88%). (D) The resistance-associated RplX A14T was absent in gonococci, however present in 83% of the commensal *Neisseria* within this sample.

Post-doxycycline selection, we also uncovered the V57M substitution in RpsJ. In natural *Neisseria* populations there were 1648 isolates harboring a valine (V) and 464 harboring a methionine (M) (Figure 2). Most of the methionine substitutions were in *N. gonorrhoeae*, with only four commensal isolates harboring a methionine at position 57: Three *N. lactamica* (88796 (isolated in Brazil in 2014), 88816 (isolated in Brazil in 2014), and 122532 (isolated in the UK in 2019)), and one in *N. subflava* (44615 isolated in the UK in 2015). One clade of *N. gonorrhoeae* isolates was distinct from wild-type *rpsJ* alleles and clustered independently (144397 (Norway 2023),144395 (Norway 2023), and 95638 (Ireland 2021)), however no clear commensal recombination tracts could be identified. In RplX, 795 isolates harbored an alanine (A), 1315 a threonine (T), and 2 a valine (V) (Figure 2). Gonococci all shared an alanine at this position, however commensals harbored the derived threonine mutation uncovered in our experimental evolution approach. There we limited evidence of recombination between commensals and pathogen at this locus, with a single *N. gonorrhoeae* clade; however one *N. lactamica* isolate (37009 isolated in the UK in 2015) appeared to have a more gonococcal-like allele at this locus (Figure 2).

### Short-term co-evolution of pathogenic and commensal *Neisseria* confirms rapid transfer of tetracycline and **β**-lactam resistance-harboring plasmids

Single colony picks were selected from each co-culture (n=2 per strain) for antibiotic susceptibility testing, for both doxycycline and penicillin, and whole genome sequencing to identify plasmid transfer events. In these isolates, compared to ancestral lineages, we observed elevated doxycycline MIC (≥ 12 μg/ml) for AR-0944 (n=2) and AR-0953 (n=2) (Table 2). These strains also all inherited the WHO N pConj plasmid. For AR-0944, the isolated co-culture strains also had an elevated penicillin MICs (≥ 32 μg/ml), which was correlated with the inheritance of the gonococcal p*bla* plasmid. p*bla* inheriting strains also inherited the gonococcal p*cryp* plasmid. For the two AR-0948 strains, one failed to grow after storage; and one interestingly had a lower doxycycline MIC upon characterization (2 μg/ml), despite selection on 6 μg/ml plates, and did not inherit any of the WHO N plasmids. For AR-0948.2, this possibly could indicate a plasmid loss event in the *in vitro* environment (Table 2). To confirm the co-culture strains investigated were indeed commensals and not the gonococcal WHO N strain, kraken2 was used to confirm species identity (Table 2). The AR-0944 and AR-0953 strains were positively identified as *N. cinerea* and *N. subflava*; however, the majority of reads for the AR-0948 derivative strain were assigned to *N. wadsworthii* (68%) rather than *N. canis* (10%). Thus, we checked the original AR-0948 read libraries (SRR12148950) (Fiore *et al*. 2020; Raisman *et al*. 2022; Frost *et al*. 2024) to confirm the kraken2 read distribution was similar. The read distribution was nearly identical (*N. wadsworthii* = 62% and *N. canis* = 9%); thus likely truly representative of the AR-0948 isolate, and not a contamination error. Of the two colonies picked from the AR-0945 co-culture, one failed to grow after storage, and one was identified as *N. gonorrhoeae*, and thus discarded from further analysis. *N. elongata* colonies are the most similar in appearance to *N. gonorrhoeae* out of the tested species, making commensal selection more difficult.

**Table 2.** MICs and WHON plasmid presence and absence data for *Neisseria* co-culture strains.

| Strain | Ancestral<br>doxycycline<br>MIC | Final<br>doxycycline<br>MIC | Ancestral<br>penicillin<br>MIC | Final<br>penicillin<br>MIC | WHO<br>N<br>pConj | WHO<br>N<br>pbla | WHO<br>N<br>pcryp | Species ID<br>kraken2 |
| --- | --- | --- | --- | --- | --- | --- | --- | --- |
| AR-0944.1 | 2 | 12 | 0.38 | >32 | + | + | + | *N. cinerea* |
| AR-0944.2 | 2 | 16 | 0.38 | >32 | + | + | + | *N. cinerea* |
| AR-0948.2 | 2 | 2 | 0.25 | 0.25 | - | - | - | *N. wadsworthii/N. canis* |
| AR-0953.1 | 0.38 | 64 | 1.5 | 1 | + | - | - | *N. subflava* |
| AR-0953.2 | 0.38 | 64 | 1.5 | 1.5 | + | - | - | *N. subflava* |

### Evidence for bystander selection: Doxycycline resistance in newly isolated commensal *Neisseria* isolates is associated with prior doxycycline use

Isolation of commensal bacteria from study participants (n=15) resulted in 150 single colony picks (n=10 per participant). Of these, 110 were positively identified as *Neisseria* after WGS and running the read libraries through kraken2: 76% of samples were identified as *N. subflava*, 14% *N. sicca* (now described as variants of *N. mucosa* (Diallo *et al*. 2019)), 9% as *Neisseria* but unidentifiable, and 1% as *N. elongata* (Figure 3; Supplementary Table 1). 51 (or 46%) of these had doxycycline MICs > 1 μg/mL, and are considered resistant as per the CLSI guidelines for tetracycline for *N. gonorrhoeae*. 62, or 56% of isolates, had intermediate levels of susceptibility (MICs = 0.5 to 1 μg/mL); and 21 (19%) were susceptible. 22 (20%) of isolates, all *N. subflava*, had MICs > 8 μg/mL. These were isolated from six study participants, with the majority originating from only two study participants with 9 (Participant 4) and 8 (Participant 15) high-level doxycycline resistant commensals isolates. Participant 4, a 36-year-old female, self-reported to have taken doxycycline in the past 6 months prior to the study; while Participant 15, a 24-year-old male, had not reported to have taken any antibiotics in the past 6 months. Of the four remaining participants with high-level doxycycline resistant commensals isolates, two did not return a questionnaire, one reported not taking antibiotics, and one had also taken doxycycline in the past 6 months (Participant 14). To assess the impact of doxycycline use in the past 6 months on doxycycline MICs, a linear model was fit followed post-hoc by Tukey’s HSD test. We found antibiotic use had a significant impact on doxycycline MIC (AOV, *p*L=L0.000069). Doxycycline MICs were significantly higher for isolates collected from participants that had used doxycycline in the last 6 months compared to participants reporting no antibiotic use (Tukey’s HSD, *p*L=L0.0002) or the group of participants that did not self-report (Tukey’s HSD, *p*L=L0.00006) (Figure 4).

**Figure 3.**
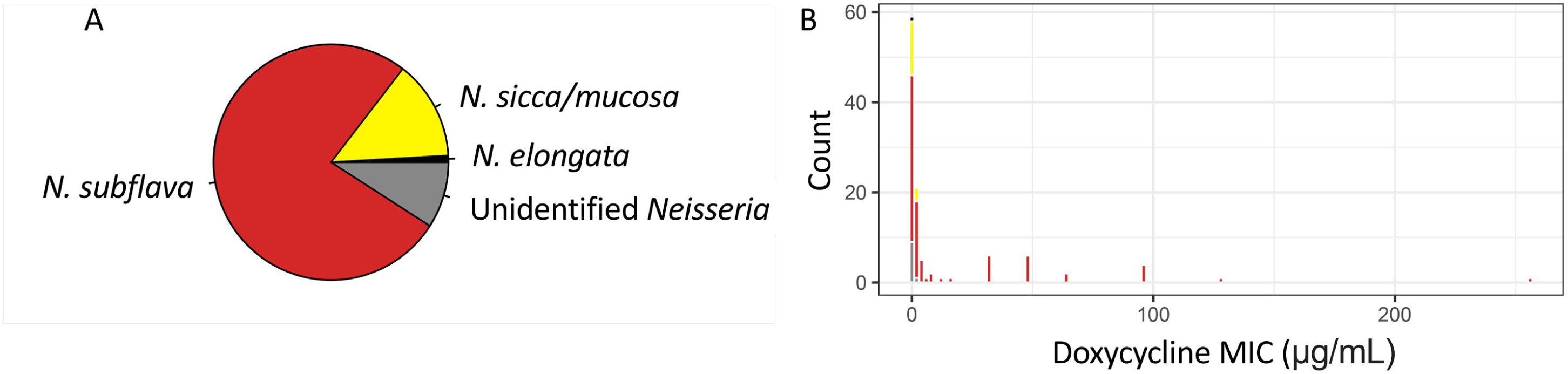
Attributes of novel commensal *Neisseria* (n=110) collected fall 2024 in Rochester, NY. (A) 76% of samples were identified as *N. subflava*, 14% *N. sicca/N. mucosa*, 9% as *Neisseria* but unidentifiable, and 1% as *N. elongata*. (B) 51 (or 46%) of isolates had doxycycline MICs > 1 μg/mL, and are considered resistant as per the CLSI guidelines for tetracycline for *N. gonorrhoeae*. 62, or 56% of isolates, had intermediate levels of susceptibility (MICs = 0.5 to 1 μg/mL); and 21 (19%) were susceptible. 22 (20%) of isolates, all *N. subflava*, had MICs > 8 μg/mL.

**Figure 4.**
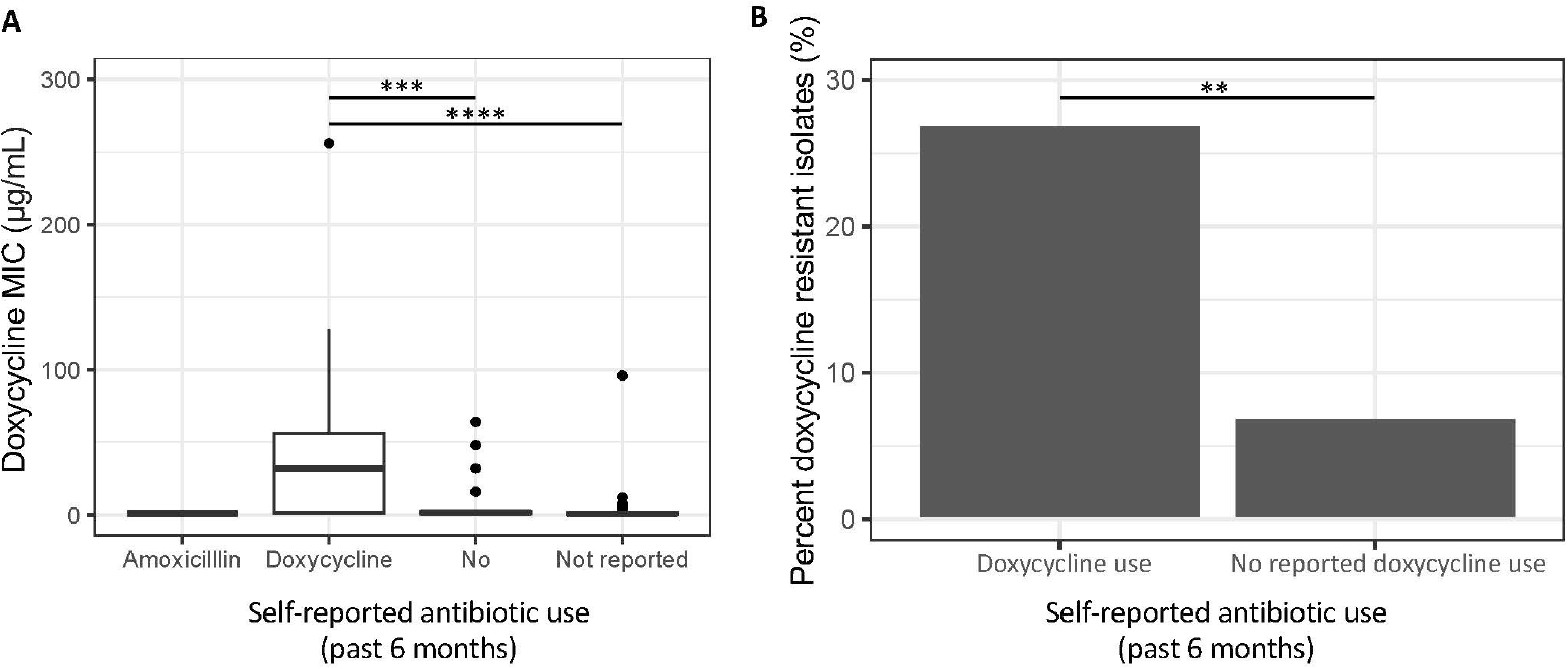
Doxycycline use is associated with doxycycline resistance in *Neisseria* commensals. Data on doxycycline use was collected from study participants at the Rochester Institute of Technology. (A) Doxycycline use in the past 6 months had a significant impact on doxycycline MIC (AOV, *p*L=L0.000069). Doxycycline MICs were significantly higher for isolates collected from participants that had used doxycycline in the last 6 months compared to participants reporting no antibiotic use (Tukey’s HSD, *p*L=L0.0002) or the group of participants that did not self-report (Tukey’s HSD, *p*L=L0.00006). (B) Additionally, there was a significant higher proportion of resistant commensals in the group that had experienced doxycycline selection in the past 6 months (27%), compared to the group that had not experienced selection (7%) (*X2 =* 8.54, *p =* 0.003).

Putative and known resistance-associated loci were characterized in these novel isolates. Here we found: Only wild-type 57V at RspJ and 46A at MtrR alleles in RIT commensals (Supplementary Table 1). However, we found a high frequency (100%) of the resistance-associated RplX A14T in these new isolates. 100% of these isolates with doxycycline MICs > 8 μg/mL carried pConj and the *tetM* gene (Supplementary Table 1). Finally, we found one *N. subflava* isolate carrying a partial gonococcal TEM sequence (Participant 10, isolate 4: 100% sequence identity to gonococcal TEM-type genes; Supplementary Table 1).

## Discussion

Doxy-PEP, or prescribing doxycycline within 24–72 h of unprotected sex as a post-exposure prophylactic therapy for STIs, has had some success in the prevention of chlamydia and syphilis (Molina *et al*. 2018; Luetkemeyer *et al*. 2023b); however, there is concern among the members of the *Neisseria* research community that long-term doxy-PEP will select for doxycycline resistant and multidrug resistant gonococci (Kong, Kenyon and Unemo 2023; Reichert and Grad 2024; Vanbaelen, Manoharan-Basil and Kenyon 2024); with mathematical modeling supporting this concern (Reichert and Grad 2024). Furthermore, doxy-PEP may have consequences for non-target commensal *Neisseria* species through bystander selection; and selecting for a greater pool of resistance in commensals is concerning given the widespread allele and gene sharing between *Neisseria* species. Limited studies have investigated doxycycline resistance in *Neisseria* commensals; a single *in vitro* study finding no evidence of resistance emergence in a 7-day span (for *N. subflava* and the pathogen *N. gonorrhoeae*) (Kenyon *et al*. 2023); and one study showing no significant increase (though a marginal one, 63 to 69% resistant isolates before and after doxycycline use respectively) in commensal doxycycline resistance rates after doxy-PEP implementation (Luetkemeyer *et al*. 2023a). Here, we reevaluate the potential for doxycycline resistance emergence in the commensal *Neisseria* after doxycycline exposure using a multifaceted approach; including *in vitro* selection, *in vitro* multi-species co-cultures, and natural population profiling of existing and novel commensal *Neisseria* isolates.

In gonococci, the main chromosomal mutation imparting reduced susceptibility is a RpsJ (ribosomal S10 protein) V57M substitution. This mutation is located proximal to the aminoacyl-tRNA site in the 30S ribosomal subunit, and alters the structure near the tetracycline-binding site, decreasing antibiotic affinity (Hu *et al*. 2005; Unemo and Shafer 2011). Therefore, it is no surprise that this mutation emerged post-doxycycline selection in an *N. elongata* with an elevated MIC compared to the ancestral lineage (0.5 to 1 μg/mL; Table 1). Additional ribosome-associated mutations also emerged (RplX A14T and *16S rRNA G1057C*) in multiple lineages of *N. subflava*, which also appeared to be associated with higher doxycycline MICs (Table 1). Finally, a MtrR A46T mutation also emerged post-selection in two lineages of *N. cinerea* with elevated MICs (from 2 to 4 μg/mL; Table 1); proximal to residue 45 which is important in mediating binding of MtrR to the promoter of the *mtrCDE efflux pump*, and modulating pump expression. These two residues sit in the helix–turn–helix (HTH) DNA binding motif of MtrR, with the G45D mutation decreasing MtrR binding to the target DNA and upregulating the pump (Shafer *et al*. 1995; Warner, Shafer and Jerse 2008). Mutations increasing *mtr* expression have been implicated in high-level tetracycline resistance previously (Hu *et al*. 2005), and therefore, though MtrR A46T has not been previously associated with tetracycline or doxycycline resistance to our knowledge, is a good candidate for further investigation. Additional doxycycline resistant lineages emerged post-selection (Table 1), however the mechanisms of resistance in these isolates were unclear at this point. Finally, the *N. subflava* lineages evolving elevated doxycycline MICs also evolved PilB reading frame altering mutations (Table 1). PilB has been shown in gonococci to be involved in survival to agents that can generate reactive oxygen species (ROS) (Skaar *et al*. 2002); and while these mutations may also contribute to elevated doxycycline MICs, the mechanism by which these would act is currently unclear as they likely produce non-functional proteins.

In the sample of natural *Neisseria* isolates from PubMLST (n=2116) (Jolley, Bray and Maiden 2018), the derived RpsJ V57M was rare in commensal isolates (∼0.3%). We found this mutation in three *N. lactamica* isolates: two isolated from Brazil in 2014 (88796 and 88816), and one isolated in the UK in 2019 (122532); and one *N. subflava* isolate from the UK in 2015 (44615). RpsJ 57M was much more prevalent in gonococci (88%). The derived MtrR A46T substitution was also more prevalent in gonococci (17%) than commensals (0.1%). Conversely however, the resistance-associated RplX A14T was absent in gonococci, however present in 83% of the commensal *Neisseria* within this sample. Though limited in scope, it is somewhat interesting to find this seemingly inverse relationship between species designation (pathogen *vs.* commensal) and presence of doxycycline-resistance associated mutations. Limited availability of mutations in particular *Neisseria* (regardless if they are causal to resistance) could suggest one of two possibilities: (i) they are maladapted to certain genomic backgrounds, and are deleterious and selected against, leaving limited opportunities for cross-species transfer, or (ii) ecological speciation and limited admixture has provided enough of a barrier to gene exchange that these alleles have not yet had the opportunity to transfer and circulate in divergent *Neisseria* clades. Regardless, it is interesting to have found experimentally derived mutations in natural *Neisseria* populations; and suggests their wide-spread availability to the *Neisseria*. Causality of these mutations in imparting resistance to doxycycline will ultimately be important to prove, which will be the subject of future investigations by this lab group.

Missing from the PubMLST dataset are reported doxycycline and/or tetracycline MICs for the majority of isolates investigated, which makes the association of genotypes to resistance phenotypes difficult. Thus, we collected a new panel of doxycycline-MIC phenotyped commensal isolates (n=110), and used WGS to identify their underlying genotypes. 51 (or 46%) of these new isolates collected from faculty, students, and staff at the Rochester Institute of Technology had doxycycline MICs > 1 μg/mL. Though seemingly high, a prior report also found high prevalence of doxycycline resistance in commensals (∼63%) collected from participants in high-risk populations for STIs (MSM/TGW living with HIV or on PrEP) (Luetkemeyer *et al*. 2023a). In this prior report, doxy-PEP implementation was not significantly shown to be associated with doxycycline resistance emergence in commensals, though a slight increase in prevalence was noted (63% resistant at baseline compared to 69% at month 12 post doxy-PEP, *p* = 0.11) (Luetkemeyer *et al*. 2023a). However, this study points to the selective potential of doxycycline on driving resistance emergence in commensals, and our results further support a clear association between doxycycline use and resistance evolution (Figure 4), with a recently published meta-analysis also supporting this finding (Vanbaelen, Manoharan-Basil and Kenyon 2024). There was a significantly higher rate of resistant commensals in the group that had experienced doxycycline selection in the past 6 months (27%), compared to the group that had not experienced selection (7%) (*X2 =* 8.54*, p =* 0.003; Figure 4). Additionally, doxycycline MICs were significantly higher for isolates collected from participants that had used doxycycline in the last 6 months compared to participants reporting no antibiotic use (Tukey’s HSD, *p*L=L0.0002) or the group of participants that did not self-report (Tukey’s HSD, *p*L=L0.00006) (Figure 4). Though our sample size is small, this may provide additional support for bystander selection of commensal doxycycline resistance after doxycycline use; which may potentially be a long-term outcome of doxy-PEP implementation.

We also searched for derived mutations from our experimental evolution study in novel commensal isolates from RIT, to see if any were associated with elevated doxycycline MICs. Interestingly, the derived RpsJ V57M was absent in RIT commensal isolates (Supplementary Table 1). This mutation is the main chromosomal mutation contributing to tetracycline-class antibiotic resistance in gonococci (Mortimer *et al*. 2022); but was absent or in very low frequency (∼0.3% in PubMLST isolates) in all natural commensal datasets characterized within this study. Similarly to the PubMLST dataset, the resistance-associated RplX A14T was present at a high frequency (100%) in the sample of commensal *Neisseria* collected from RIT. Finally, also mirroring the PubMLST dataset, the derived MtrR A46T was absent in the RIT commensal panel (present in only 0.1% of PubMLST isolates). For commensal *N. subflava* isolates with high doxycycline MICs (> 8 μg/mL), 100% of them carried the *tetM* gene (> 97% sequence identity to the WHO N gonococcal *tetM* gene; Supplementary Table 1). Fiore et al. (2020) also characterized a *N. subflava* isolate (AR-0954) with a high tetracycline MIC (48 μg/mL) carrying *tetM*. There was one isolate unidentified *Neisseria* isolate collected from Participant 14 however, which also appeared to carry *tetM* (WHO N *tetM* % identity: 92%), yet had a lower-than-expected doxycycline MIC (4 μg/mL). It is unclear at this point why their doxycycline MICs were low, despite carriage of *tetM*. In gonococci, there are three main variants of the β-lactamase gene on p*bla* plasmids, TEM-1 (allele 3) present in ∼60% of gonococci, TEM-135 (allele 2) present in 24% of isolates, and TEM-1 with a P14S substitution (allele 6) found in 11% of isolates (Elsener *et al*. 2023). TEM-135 carries M182T, located in the hinge region between two β-lactamase domains, which aids in the stability of the enzyme compared to TEM-1 (Orencia *et al*. 2001; Kather *et al*. 2008; Salverda, De Visser and Barlow 2010). Here, we find one *N. subflava* isolate carrying a gonococcal TEM gene (Participant 10, isolate 4: Supplementary Table 1). The TEM gene in this *N. subflava* isolate was a partial hit to gonococcal TEM-1 (allele 3) and TEM-135 (allele 2) alleles, having 100% similarity to both at nucleotide positions 619 to 891 (252 bps) (Supplementary Figure 1). This partial hit was present on a single contig in the isolate’s assembly (Contig 2584), which had a total length of 252 bps. Unfortunately, this sequence did not include sites which would help to identify the TEM allele present within this isolate.

The pConj plasmid will undoubtedly increase in prevalence in the Dox-PEP era, and as it contains all of the necessary machinery for conjugation, will likely facilitate spread of the p*bla* plasmid and β-lactam resistance across *Neisseria* populations. Multiple studies support this idea, including those that show associations of pConj and p*bla* carriage in natural *N. gonorrhoeae* populations (Cehovin *et al*. 2020), and laboratory experiments that show cross-species transfer of both plasmids between *N. gonorrhoeae* and multiple commensal species (Genco, Knapp and Clark 1984; Roberts and Knapp 1988; Roberts and Knapp, JS 1988). Here, we provide further support for this idea using a different experimental framework. The gonococcal WHO N strain was cultured with *N. cinerea*, *N. elongata*, *N. canis*, and *N. subflava* for 7 days in the absence of selection, and on the final day strains were plated on doxycycline plates and single colonies were picked. In this context we found pConj transfer to *N. cinerea* and *N. subflava*, in addition to p*bla* transfer to *N. cinerea*. Interestingly, a *N. canis* strain in our experiment appeared to have acquired pConj (i.e., as evidenced by its transient ability to survive on 6 μg/mL) however may have lost the plasmid after passaging. Genco et al. (1984) also note limited stability of gonococcal plasmids in some commensal strains, suggesting that some species may be more likely reservoirs of these important mobile genetic elements than others. Here, we reemphasize and highlight the idea that commensal *Neisseria* may serve as important reservoirs of plasmid-borne doxycycline and β-lactam resistance, which may become mobilized in the age of Dox-PEP. Importantly, with some variants of p*bla* only 1-2 mutations away from becoming extended-spectrum beta-lactamases (ESBLs) which can hydrolyze both penicillins and cephalosporins (i.e., *bla*_TEM-135_, requires G238S or E240K, or *bla*_TEM-1_, requires M182T and G238S or E240K) (Salverda, De Visser and Barlow 2010; Muhammad *et al*. 2014; Kandinov *et al*. 2022; Elsener *et al*. 2023), a nightmare scenario for retaining our ability to treat gonococcal infections with our last-line drug ceftriaxone, selective pressures promoting the continued spread and evolution of these plasmids such as doxy-PEP should be a major concern for public health policy makers.

In conclusion, our work underscores the importance of commensal *Neisseria* as reservoirs of doxycycline resistance for the commensal *Neisseria*. We find both target (RspJ, RplX, 16S rRNA) and efflux mutations (MtrR) as possible chromosomal contributors to resistance, however ultimately transformation experiments into ancestral backgrounds will be necessary to confirm the causality of candidate chromosomal resistance loci nominated in this study, which will be the focus of future research initiatives. Though we find limited evidence of HGT at these loci between commensals and pathogens in *Neisseria* populations, we do find evidence of some of our experimentally derived mutations circulating in these natural populations (RpsJ V57M, RplX A14T, and MtrR A46T). Our work also demonstrates a link between recent doxycycline use and the emergence of doxycycline resistance in commensal *Neisseria*, supporting prior findings by other groups (Luetkemeyer *et al*. 2023a; Vanbaelen, Manoharan-Basil and Kenyon 2024), and highlighting the very real reality and danger of bystander selection. doxy-PEP implementation thus should be a concern for driving resistance emergence in commensal communities, which in the novel commensal strains isolated herein, appears to be mainly associated with acquisition of the pConj plasmid and the *tetM* gene. A secondary danger to pConj acquisition, is spread of p*bla* and β-lactam resistance, which we demonstrate here *in vitro*, and has been documented before (Genco, Knapp and Clark 1984; Roberts and Knapp 1988; Roberts and Knapp, JS 1988). Thus, we believe careful monitoring of both gonococcal and commensal *Neisseria* populations should be considered in communities implementing doxy-PEP, especially surveillance of plasmid carriage, to ensure high-level doxycycline and β-lactam resistance is not increasing.

## Data Availability

All scripts and datasets are available on:(Knapp 1988; Knapp and Hook 1988)https://github.com/wadsworthlab. Read libraries generated in this study can be accessed on the Sequence Read Archive: Doxycycline selected strains (BioProject: PRJNA1018855), co-evolved strains (PRJNA1207413), and novel commensal strains (BioProject: PRJNA1208441).

## Acknowledgements

The authors would like to acknowledge the generous support provided by the RIT College of Science and the Thomas H. Gosnell School of Life Science for this work. Work reported in this publication was also supported by the National Institute of General Medical Sciences of the National Institutes of Health under Award Number R15AI174182. The content is solely the responsibility of the authors and does not necessarily represent the official views of the National Institutes of Health. The funders had no role in study design, data collection and analysis, decision to publish, or preparation of the manuscript. The authors would also like to thank Girish Kumar at the RIT Genomics Core for providing support and sequencing services.

---

**Supplementary Figure 1.** Alignment of the p*bla* β-lactamase gene variants TEM-1 and TEM-135. Here, we identify one novel *N. subflava* isolate carrying a TEM gene (Participant 10, isolate 4). The TEM gene in isolate was a partial hit to TEM-1 (allele 3) and TEM-135 (allele 2) alleles, having 100% similarity to both at nucleotide positions 619 to 891 (252 bps). This partial hit was present on a single contig in the isolate’s assembly (Contig 2584), which had a total length of 252 bps. This partial sequence did not include sites (i.e., 182) which would allow identification of the TEM allele present within this isolate.

