## Supplementary figures and images for "Re-visiting the potential impact of doxycycline post-exposure prophylaxis (doxy-PEP) on the selection of doxycycline resistance in *Neisseria* commensals"

### Supplementary Figure 1

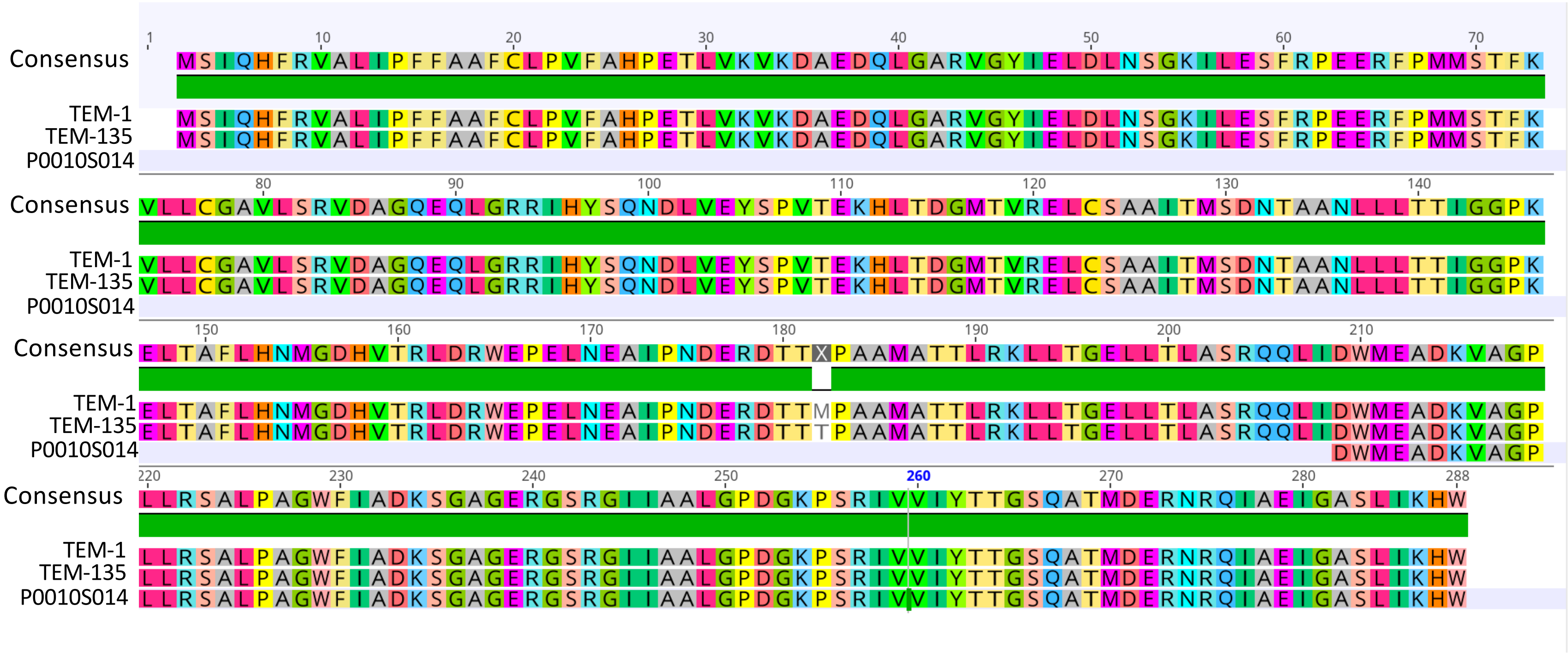
